# α-Synuclein-112 impairs synaptic vesicle recycling consistent with its enhanced membrane binding properties

**DOI:** 10.1101/2020.04.03.024125

**Authors:** Lindsey G. Soll, Julia N. Eisen, Karina J. Vargas, Audrey T. Medeiros, Katherine M. Hammar, Jennifer R. Morgan

## Abstract

Synucleinopathies are neurological disorders associated with α-synuclein overexpression and aggregation. While it is well established that overexpression of wild type α-synuclein (α-syn-140) leads to cellular toxicity and neurodegeneration, much less is known about other naturally occurring α-synuclein splice isoforms. In this study we provide the first detailed examination of the synaptic effects caused by one of these splice isoforms, α-synuclein-112 (α-syn-112). α-Syn-112 is produced by an in-frame excision of exon 5, resulting in deletion of amino acids 103-130 in the C-terminal region. α-Syn-112 is upregulated in the substantia nigra, frontal cortex, and cerebellum of parkinsonian brains and is correlated with susceptibility to sporadic Parkinson’s disease (PD), dementia with Lewy bodies (DLB) and multiple systems atrophy (MSA). We report here that α-syn-112 binds strongly to anionic phospholipids when presented in highly-curved liposomes, similar to α-syn-140. However, α-syn-112 bound significantly stronger to all phospholipids tested, including the phosphoinositides. α-Syn-112 also dimerized and trimerized on isolated synaptic membranes, while α-syn-140 remained largely monomeric. When introduced acutely to lamprey synapses, α-syn-112 robustly inhibited synaptic vesicle recycling. Interestingly, α-syn-112 produced effects on the plasma membrane and clathrin-mediated synaptic vesicle endocytosis that were phenotypically intermediate between those caused by monomeric and dimeric α-syn-140. These findings indicate that α-syn-112 exhibits enhanced phospholipid binding and oligomerization *in vitro* and consequently interferes with synaptic vesicle recycling *in vivo* in ways that are consistent with its biochemical properties. This study provides additional evidence suggesting that impaired vesicle endocytosis is a cellular target of excess α-synuclein and advances our understanding of potential mechanisms underlying disease pathogenesis in the synucleinopathies.

## INTRODUCTION

Synucleinopathies are a class of neurological disorders linked to overexpression and aggregation of α-synuclein, and they include Parkinson’s Disease (PD), Dementia with Lewy Bodies (DLB), and Multiple Systems Atrophy (MSA). In these diseases, α-synuclein aggregates throughout neurons, including axons and synapses, leading to cellular toxicity and neurodegeneration (Kramer and Schulz-Schaeffer, 2007; Schulz-Schaeffer, 2010; Scott et al., 2010; Burre et al., 2018; Sulzer and Edwards, 2019). Gene multiplication and an increasing number of point mutations in the α-synuclein gene *(SNCA)* are known to lead to α-synuclein aggregation and are associated with both inherited and sporadic forms of PD, DLB, and various forms of parkinsonism (Kruger et al., 1998; Singleton et al., 2003; Nussbaum, 2018). Thus, it is increasingly important to understand how different α-synuclein variants impact neuronal function, as well as disease pathogenesis and progression.

The wild type α-synuclein gene, *SNCA,* comprises six exons, which translates into a protein of 140 amino acids. To date, four additional splice isoforms have been identified: α-syn-126, α-syn-112, α-syn-98, and α-syn-41. α-Syn-126, α-syn-112, and α-syn-98 comprise in-frame deletions of exon 3, exon 5, or both, respectively, resulting in shorter protein products (Ueda et al., 1994; Beyer et al., 2008; McLean et al., 2012). α-Syn-41 lacks exons 3 and 4, generating an early stop codon and resulting in a truncated N-terminal peptide (Vinnakota et al., 2018). All of these splice isoforms are expressed in control brains and exhibit differential expression in PD, DLB, and Alzheimer’s disease, with levels generally being higher in the diseased brains (Beyer et al., 2004; Beyer et al., 2008; McLean et al., 2012; Cardo et al., 2014). In comparison to wild type α-syn-140, surprisingly little is known about its splice isoforms and how they affect neuronal functions.

In this study, we focus on the splice isoform α-syn-112, a 12 kDa protein comprising a deletion of 28 amino acids (a.a. 103-130) near the C-terminus (Ueda et al., 1994). α-Syn-112 is normally expressed in low levels in many human tissues, including skin, lung, kidney, and heart, with highest expression in the brain (Beyer et al., 2008). However, in parkinsonian, DLB and MSA brains, α-syn-112 is overexpressed in the substantia nigra, frontal cortex, and cerebellum (Beyer et al., 2004; Brudek et al., 2016). In addition, increased α-syn-112 levels are associated with PD risk (McCarthy et al., 2011). Compared to α-syn-140, α-syn-112 exhibits enhanced aggregation and fibrillation *in vitro* (Manda et al., 2014). While it is clear that excess α-syn-112 is associated with a number of neurodegenerative diseases, very little is known about its biochemical properties or neuronal functions.

We therefore set out to perform a more detailed characterization of α-syn-112, focusing on its possible roles at synapses. Under physiological conditions, α-syn-140 is expressed at the presynapse where it regulates synaptic vesicle clustering and trafficking (Bendor et al., 2013; Vargas et al., 2014; Logan et al., 2017; Atias et al., 2019). When overexpressed at mammalian synapses to levels comparable to those in familial PD, α-syn-140 impaired synaptic vesicle trafficking (Nemani et al., 2010; Scott et al., 2010), and altered the composition of presynaptic proteins (Scott et al., 2010). In line with these findings, we previously reported that acute introduction of α-syn-140 at a classical vertebrate synapse, the lamprey reticulospinal (RS) synapse, impaired synaptic vesicle recycling in ways consistent with deficits in clathrin-mediated endocytosis and possibly bulk endocytosis (Busch et al., 2014; Medeiros et al., 2017; Banks et al., 2020). Similarly, acute introduction of α-syn-140 at mammalian synapses also impaired vesicle endocytosis with no observable effects on exocytosis (Xu et al., 2016; Eguchi et al., 2017). The synaptic deficits induced by α-syn-140 require proper membrane binding because point mutants with reduced lipid binding capacity exhibited greatly reduced effects on SV trafficking (Nemani et al., 2010; Busch et al., 2014). In comparison, there are no studies to date that have investigated how any of the related α-synuclein splice isoforms affect presynaptic functions, prompting this work.

Here we describe the membrane binding properties of α-syn-112 and its corresponding effects at synapses. It is well established that α-syn-140 binds to anionic phospholipids, such as phosphatidic acid (PA) and phosphatidylserine (PS), especially when presented in small, highly curved liposomes (Davidson et al., 1998; Burre et al., 2010; Burre et al., 2012; Busch et al., 2014). In comparison to α-syn-140, α-syn-112 bound more strongly *in vitro* to all phospholipids tested, including phosphoinositides that regulate synaptic vesicle trafficking such as PI(4)P and PI(4,5)P_2_ (Di Paolo and De Camilli, 2006; Saheki and De Camilli, 2012). In addition, α-syn-112 had a greater propensity for oligomerization on purified synaptic membranes. Consistent with enhanced membrane binding and oligomerization, α-syn-112 inhibited synaptic vesicle recycling at lamprey synapses and produced a phenotype that was intermediate between monomeric and dimeric α-syn-140 (Busch et al., 2014; Medeiros et al., 2017; Medeiros et al., 2018; Banks et al., 2020). These findings implicate α-syn-112 in inducing defective synaptic vesicle trafficking, which may lead to cellular toxicity in the synucleinopathies.

## MATERIALS AND METHODS

### SDS-PAGE and Western Blotting

Recombinant human α-syn-140 and α-syn-112 were purchased from rPeptide (Bogart GA). Proteins were run on a 12% SDS-PAGE gel and then stained with Coomassie or transferred to nitrocellulose for Western blotting. For Coomassie gels, 2-3 μg of protein was loaded per lane. For Western blots, 0.2-0.3 μg of protein, or 20 μL of liposome binding assay samples, were loaded. Western blots were performed using standard procedures (Busch et al., 2014). After blocking in TBST buffer (20 mM Tris pH 7.6, 150 mM NaCl, 0.1% Tween 20) with 1% dry milk, the membranes were incubated for 2 hrs with a rabbit polyclonal pan-synuclein antibody (1:1000; ab53726; Abcam, Cambridge, MA). After washing in TBST, the blots were incubated for 1 hr with goat anti-rabbit HRP conjugated IgG (H+L) (1:1000; Thermo Scientific, Waltham, MA). Pierce^TM^ ECL Western blotting substrate (Thermo Scientific, Waltham, MA) was used to develop blots.

### Liposome Binding Assays

The lipids acquired from Avanti Polar Lipids, Inc. (Alabaster, AL) included 16:0-18:1 phosphatidylcholine (PC), 18:1-12:0 nitrobenzoxadiazole-PC (NBD-PC), 16:0-18:1 1-phosphatidic acid (PA), 16:0-18:1 phosphoethanolamine (PE), and porcine brain total lipid extract (BTLE). Phosphoinositides were acquired from Echelon Biosciences, Inc. (Salt Lake City, UT): phosphoinositol diC_16_ (PI), phosphoinositol 3-phosphate diC_16_ [PI(3)P], phosphoinositol 4-phosphate diC_16_ [PI(4)P], phosphoinositol 5-phosphate diC_16_ [PI(5)P], phosphoinositol 3,4-bisphosphate diC_16_ [PI(3,4)P_2_], phosphoinositol 3,5-bisphosphate diC_16_ [PI(3,5)P_2_], phosphoinositol 4,5-bisphosphate diC_16_ [PI(4,5)P_2_], and phosphoinositol 3,4,5-triphosphate diC_16_ [PI(3,4,5)P_3_].

Liposome binding assays were performed as previously described (Burre et al., 2012; Busch et al., 2014; Medeiros et al., 2017). Artificial liposomes were generated by mixing lipids (1.0 mg total) in the desired proportions in 200 μL of 2:1 chloroform:methanol and then dried into a monolayer by a stream of N_2_. All liposomes contained 1% fluorescently labeled NBD-PC. After drying, sucrose (1 mL; 300 mM) was added, and lipids were incubated at 37°C for 20 min to swell the membranes, followed by vortexing for 1 min to generate large, heterogeneous liposomes. Small unilamellar vesicles (SUVs; 30-50 nm) were generated by probe sonication for 5 sec at 15 sec intervals for 2 min at RT. Liposomes were then centrifuged at 80,000 x g for 20 minutes at RT and supernatant, containing the SUVs, was isolated for the liposome binding assay.

α-Syn140 or α-syn112 (5 mg) was incubated for 2 hrs at RT with liposomes (~34-36 μL), in HKE buffer (25 mM HEPES, pH 7.4, 150 mM KCl, 1 mM EDTA) up to 100 μL total volume. Samples were then added to the bottom of an Accudenz gradient (40%, 35%, 30%, and then 0%, 800 μL total volume). Columns were centrifuged at 280,000 x g for 3 hrs at RT. After ultracentrifugation, columns were separated into 8 fractions (100 μL each). The presence of liposomes was determined in each fraction by quantifying NBD fluorescence using a Nanodrop 3300. In parallel, the α-synuclein distribution in each fraction was determined by Western blotting. Quantification of α-synuclein band intensities was performed using ImageJ. Lipid bound protein (%) was calculated as the amount of protein in the first 3 fractions divided by the total protein in all 8 fractions. Data shown are representative of n=3-7 independent experiments. Graphpad Prism 8 (GraphPad Software, Inc., La Jolla, CA) was used to perform statistical analyses and generate graphs.

### Membrane Recruitment Assays

Membrane recruitment assays were performed as described (Shetty et al., 2013). First, crude synaptosomes were prepared from mouse brain, resuspended in 6 mL homogenization buffer (25 mM Tris-HCl; pH 8.0, 500 mM KCl, 250 mM sucrose, 2 mM EGTA), then added to the top of freshly prepared 0.65 M, 0.85 M, 1.00 M, 1.20 M sucrose gradients and centrifuged at 100,000 x g for 2 hrs. Synaptosomes were collected from the 1/1.2 M interface and resuspended in 20 mL buffer. Pure synaptosomes were centrifuged at 100,000 x g for 20 min. The pellet was resuspended in 4 mL icecold deionized water, and 250 mM HEPES-NaOH, pH 7.4 was added to a final concentration of 7.5 mM. The suspension was incubated on ice for 30 min and centrifuged at 100,000 x g for 20 min. The pellet was resuspended in 4 mL of 0.1 M Na_2_CO_3_ to strip peripheral proteins, incubated for 15 min at 37°C, and centrifuged at 100,000 x g for 20 min. Pellet was resuspended in 2 mL cytosolic buffer (25 mM HEPES-NaOH, pH 7.4, 120 mM potassium glutamate, 2.5 mM magnesium acetate, 20 mM KCl, and 5 mM EGTA-NaOH, pH 8.0, filtered and stored at 4°C), centrifuged again at 100,000 x g for 20 min, and resuspended in 2 mL of cytosolic buffer. Proteins were quantified using BCA. Mini cOmplete^TM^ protease inhibitors (Roche) were added, and aliquots of purified membrane resuspension were flash frozen and stored at −80°C until use.

Next, cytosol preparations were made from two mouse brains. To do that, brains were first washed and then homogenized with 2 mL of homogenization buffer (25 mM Tris-HCl, pH 8.0, 500 mM KCl, 250 mM sucrose, 2 mM EGTA, and 1 mM DTT) using 10 strokes at 5,000 rpm. The homogenate was transferred to a 3.5 mL ultracentrifuge tube and centrifuged at 160,000 x g for 2 hrs at 4°C. The supernatant was exchanged into 3.5 mL cytosolic buffer. After measuring protein concentration and adding protease inhibitors, 100 μL aliquots were flash frozen and stored at −80°C until use.

For the membrane recruitment assays, synaptic membranes (200 μg) were mixed with 250 μg brain cytosol proteins in 500 μl cytosolic buffer and supplemented with different concentrations of recombinant human α-syn-140 and α-syn-112 (rPeptide). A control experiment was prepared with only synaptic membranes and cytosolic buffer. Mixtures were incubated at 37°C for 15 min. The samples were immediately centrifuged at 100,000 x g for 30 min at 4°C. Pellets, now containing the synaptic membranes with bound proteins, were resuspended in 500 μL of cytosolic buffer at 4°C. The resuspension was centrifuged at 100,000 x g for 30 min at 4°C and resuspended in 90 μL of cytosolic buffer. For each sample, 20 μL aliquots were mixed with 5X loading buffer, run on 12% reducing SDS-PAGE gels, transferred to nitrocellulose membranes, and processed via Western blotting. Levels of α-synuclein recruited to synaptic membranes were detected for each condition by Western blot with a rabbit polyclonal pan-synuclein antibody (1:1000; Abcam ab53726; Cambridge, MA) and quantified using ImageJ.

### Microinjections and Stimulation

Recombinant human α-syn-140 and α-syn-112 were obtained from rPeptide, Inc. The recombinant α-syn-140 dimer (NC dimer) used in this study was a single polypeptide comprising two full-length copies of α-syn-140, as previously described (Pivato et al., 2012; Medeiros et al., 2017). All animal procedures were conducted in accordance with standards set by the National Institutes of Health and approved by the Institutional Animal Care and Use Committee at the Marine Biological Laboratory. Tricaine-S (MS-222; 0.1 g/L; Western Chemical Inc. Ferndale, WA) was used to anesthetize late larval lampreys *(Petromyzon marinus;* 11-13 cm; M/F). Next, spinal cords were dissected into 2-3 cm segments, pinned in a Sylgard lined dish, and prepared for microinjection, as previously described (Morgan et al., 2004; Busch et al., 2014; Walsh et al., 2018). Using a glass microelectrode, α-syn140 or α-syn112 (120-200 μM) dialyzed in lamprey internal solution (180 mM KCl, 10mM HEPES K^+^, pH 7.4) was injected into reticulospinal axons using small pulses of N_2_ (4-20 ms, 30-50 psi, 0.2 Hz), which were delivered using a Toohey spritzer. We estimate that proteins were diluted 10-20x in the axon based on the fluorescence of a coinjected dye (fluorescein dextran 100 μM; 10kDa), resulting in a final axonal concentration of 7-16 μM. This concentration is 2-5 times above the estimated endogenous levels and within range of overexpression levels for animal models of PD (Nemani et al., 2010; Scott et al., 2010; Westphal and Chandra, 2013) and human patients (Singleton et al., 2003). After protein injection, short, depolarizing current pulses (30-50 nA; 1 ms) were delivered to the axons to stimulate action potentials (20 Hz, 5 min). Immediately following the stimulation period, spinal cords were fixed in 3% glutaraldehyde, 2% paraformaldehyde in 0.1M Na cacodylate, pH 7.4 for standard transmission electron microscopy.

### Electron Microscopy and Imaging

After fixation (overnight to 2 days), spinal cords were processed in 2% osmium, stained *en bloc* with 2% uranyl acetate, and embedded in Embed 812 resin, as previously described (Busch et al., 2014; Medeiros et al., 2017; Walsh et al., 2018). Ultra-thin sections (70 nm) were counterstained with 2% uranyl acetate followed by 0.4% lead citrate. A JEOL JEM 200CX transmission electron microscope was used to acquire images of individual synapses at 37,000X magnification. For each experimental condition, images were acquired from at least n=10-20 synapses collected from n=2 axons/animals at distances of 25-150 μm from the injection site, which is where the protein had diffused based on the co-injected fluorescent dye. Images of control synapses were acquired from the same axons, but at greater distances from the injection site (>350 μm) in regions where the injected proteins had not diffused, providing an internal control for each experiment.

A morphometric analysis was performed on all synaptic membranes within 1 μm of the active zone, as previously described (Busch et al., 2014; Medeiros et al., 2017; Walsh et al., 2018; Banks et al., 2020). Image analysis was performed in FIJI 2.0.0. by a researcher blinded to experimental conditions. Measurements included the number of synaptic vesicles per synapse (per section), size of plasma membrane (PM) evaginations, number and size of large (>100 nm) irregularly shaped intracellular membranous structures (“cisternae”), and number and stage of clathrin coated pits (CCPs) and clathrin coated vesicles (CCVs). The sizes of plasma membrane evaginations were measured by first drawing a straight line (1 μm) laterally from the edge of the active zone to the nearest point on the axolemma and then measuring the curved distance between these two points. Additionally, we also quantified the depth of the plasma membrane evaginations from the axolemmal surface to the deepest point within the evagination. CCP/V stages were defined as: stage 1 – initial clathrin coated bud; stage 2 – invaginated CCP without constricted neck; stage 3 – invaginated CCP with constricted neck; stage 4 – free CCV. GraphPad Prism 8 was used to generate graphs and for all statistical analyses.

Reconstruct software (Fiala, 2005) was used to generate a threedimensional reconstruction of single synapses from four or five serial images. Fiduciary markers were used to align the serial images. Synaptic structures were added using trace slabs for plasma membrane and cisternae, spheres for synaptic vesicles (50 nm) and clathrin-coated pits and vesicles (90 nm), and a Boissonnat surface for the active zone.

## RESULTS

### α-Syn-112 exhibits enhanced binding to liposomes containing anionic phospholipids

We began by comparing the lipid binding properties of α-syn-112 and α-syn-140. α-Syn-140 comprises an amphipathic alpha-helical region that is involved in lipid binding (a.a. 1-101); a non-amyloid component (NAC) domain that is involved in self-association (a.a. 61-95); and a less structured acidic C-terminal domain (a.a. 102-140) (Fig. 1A). It is well established that the N-terminal alpha helical domain of α-syn-140 (a.a. 1-95), which includes the NAC domain, is responsible for its strong binding to liposomes containing anionic phospholipids (Davidson et al., 1998; Chandra et al., 2003; Burre et al., 2012; Busch et al., 2014) and that this binding is modulated the C-terminus (Lautenschlager et al., 2018). In α-syn-112, the removal of exon 5 brings together amino acids 102 and 131 into a single polypeptide, generating a shorter 112-amino acid protein (Fig. 1A). Compared to the known NMR structure of folded, liposome-bound α-syn-140 (Ulmer et al., 2005), the predicted structure of α-syn-112 suggests that the deletion results in additional alpha-helical content in the C-terminal domain (Fig. 1B) (SWISS-MODEL, https://swissmodel.expasy.org/). The size difference between α-syn-140 and α-syn-112 is apparent by SDS-PAGE gel and Western blot (Fig. 1C).

**Figure 1.**
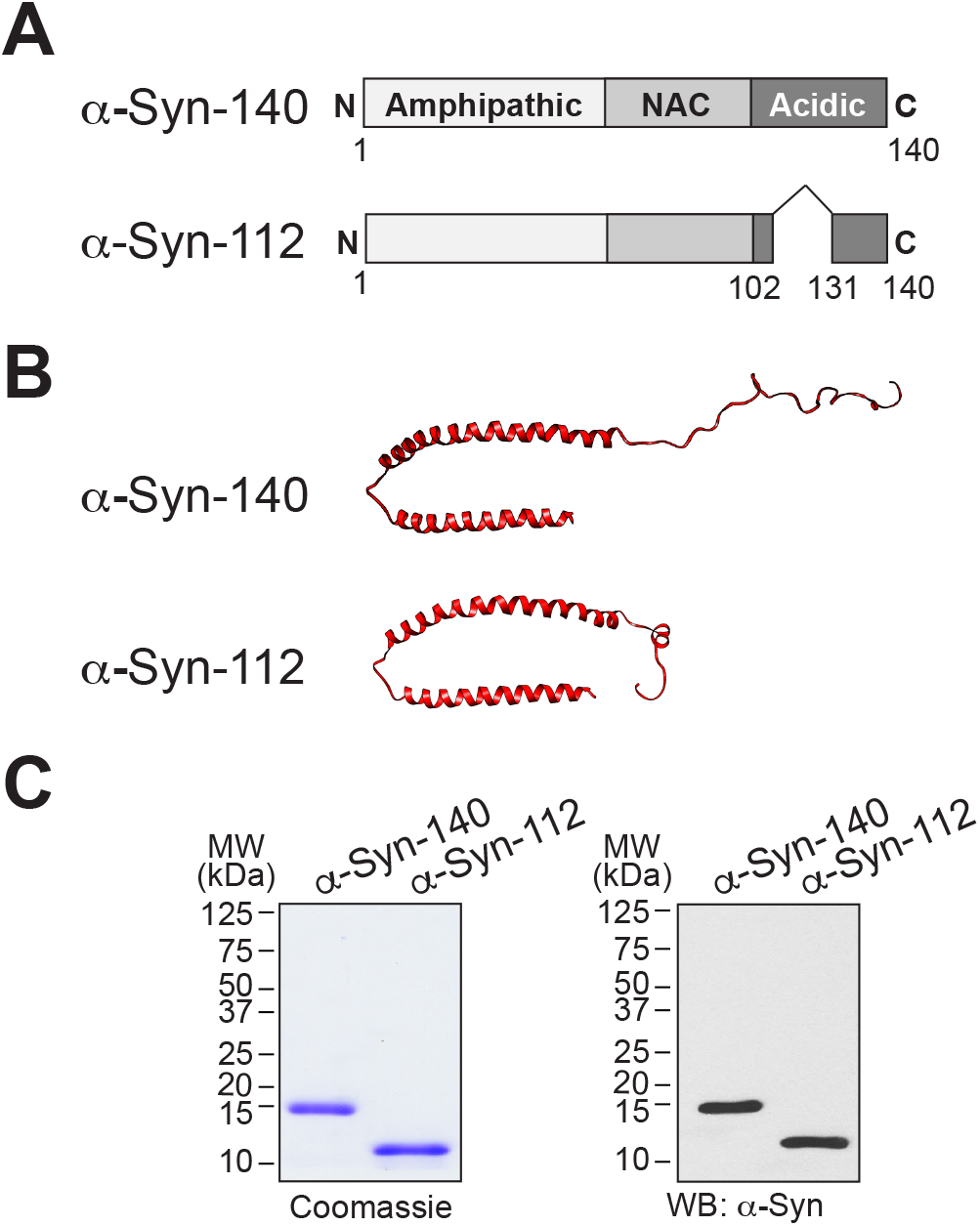
α -Syn-112 splice isoform. **A.** Domain diagram of α-syn-140 and α-syn-112 showing the result of exon 5 deletion. **B.** (top) The NMR structure of human α-syn-140 bound to lipid micelles (Ulmer et al., 2005), and (bottom) predicted structure of α-syn-112 (UCSF Chimera software). α-Syn-112 is predicted to possess a slightly extended alpha helix. **C.** Coomassie-stained SDS-PAGE gel and corresponding Western blot showing the size difference between α-syn-140 and α-syn-112.

Though it is well established that α-syn-140 binds to anionic phospholipids such as phosphatidic acid (PA) and phosphatidylserine (PS), the lipid binding properties of α-syn-112 are undetermined. We therefore used a well established liposome floatation assay to test binding of α-syn-112 to standard anionic lipids, starting with PA. The liposomes used in this assay were small unilamellar vesicles approximating the size of synaptic vesicles (30-50 nm). Under these conditions, after ultracentrifugation in an Accudenz gradient, the vast majority (>90%) of liposomes float to the top of the column in fractions 1-3, as determined by NBD fluorescence (Fig. 2A). Thus, any liposome-bound α-syn-140 also floats to the top of the column (fractions 1-3), while unbound protein remains lower in the column (fractions 4-8), as determined by Western blotting. As previously reported (Busch et al., 2014), >95% of total α-syn-140 was bound to liposomes containing 1:1 phosphatidylcholine (PC) and PA, while negligible binding was detected to liposomes containing only PC, indicating strong binding to PA (Fig. 2B-C) (PC: 4.98% ± 2.27%, n=3; PC/PA: 98.29% ± 0.72%, n=3; p<0.0001; Student’s t-test). To determine the extent to which α-syn-112 binds PA, we performed a concentration series with varying concentrations of PA, ranging from 0-50%. Surprisingly, α-syn-112 partially bound to liposomes containing only PC (0% PA), a condition in which α-syn-140 binding is negligible (Fig. 2D-E). At 0.5%, 1%, 5%, and 10% PA, α-syn-112 had enhanced liposome binding compared to α-syn-140, while no difference was detected at 50% PA where protein binding was saturated (Fig. 2D-E) [(*0% PA=* α-syn-140: 3.03 ± 1.86, n=3; α-syn-112: 34.42 ± 8.28, n=3; p<0.001) (*0.5% PA*= α-syn-140: 5.76 ± 1.89, n=4; α-syn-112: 37.697 ± 4.95, n=3; p<0.001) (*1% PA*= α-syn-140: 11.74 ± 0.93, n=3; α-syn-112: 65.08 ± 3.98, n=3; p<0.0001) (*5% PA*= α-syn-140: 28.21 ± 5.14, n=4; α-syn-112: 81.46 ± 11.32, n=3; p<0.0001) (*10% PA*= α-syn-140: 56.31 ± 5.85, n=4; α-syn-112: 93.53 ± 3.02, n=3; p=0.0002) (*50% PA*= α-syn-140 96.70 ±, n=3; α-syn-112: 92.67 ± 5.87, n=3; p=0.54 (n.s.); ANOVA Sidak’s Post Hoc]. Analysis of the binding curves revealed that α-syn-112 had a 10-fold higher affinity for PA than α-syn-140 (α-syn140: EC_50_ = 9.56% PA, R^2^=0.96; α-syn112: EC_5_0 = 0.94% PA, R^2^=0.84; Non-linear fit: dose-response). Thus, α-syn-112 exhibits much stronger binding to PC/PA liposomes than α-syn-140.

**Figure 2.**
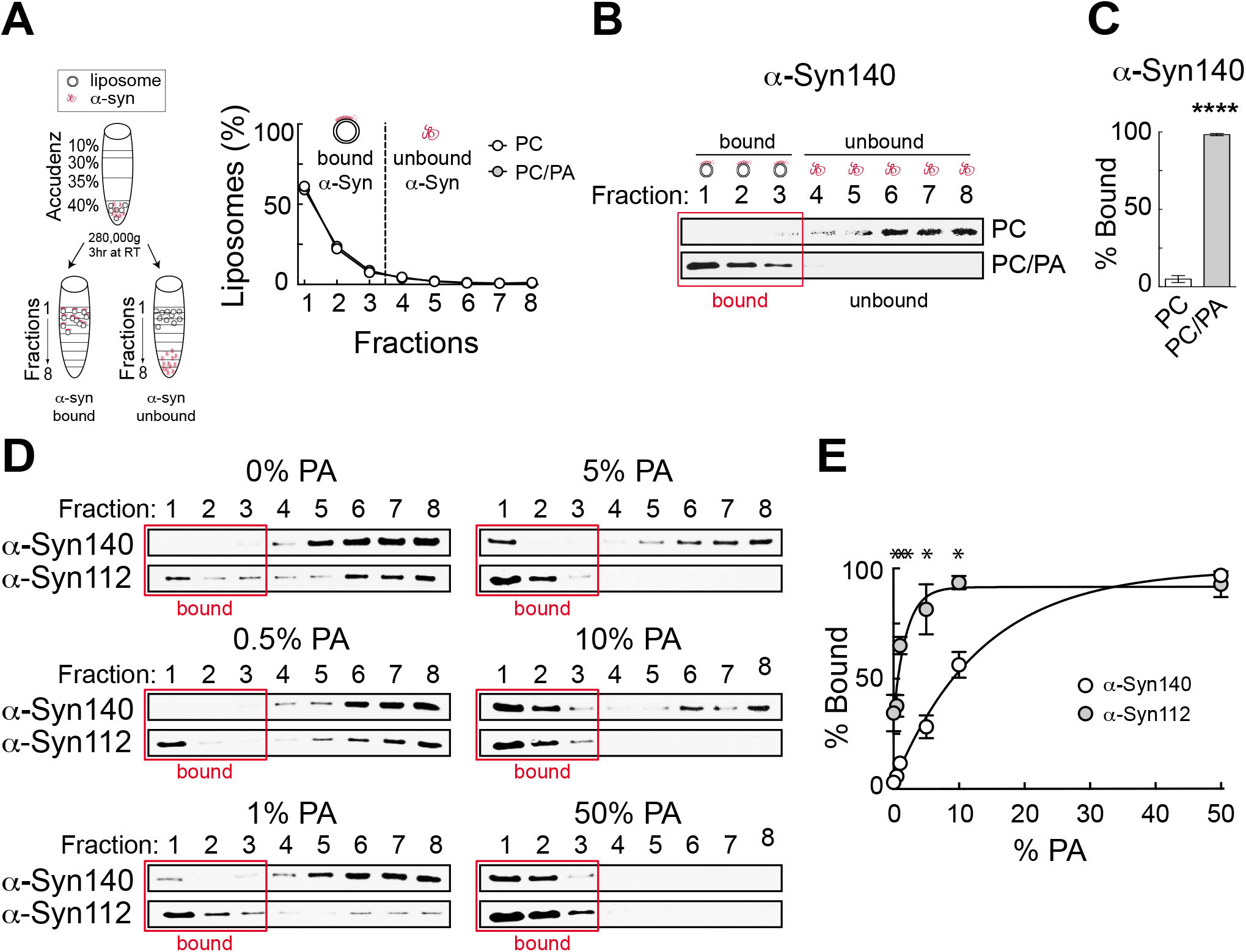
Compared to α-syn140, α-syn-112 isoform has enhanced binding to PC/PA liposomes. **A.** (left) Diagram showing the liposome binding assay. (right) After ultracentrifugation, liposomes float to the top of the gradient (fractions 1-3), carrying along any bound α-synuclein while free α-synuclein remains in the bottom fractions (4-8). **B.** Representative Western blots showing α-syn-140 binding to PC or PC/PA liposomes. α-Syn-140 binds strongly to anionic phospholipids such as PA. **C.** Quantification of lipid bound α-syn-140 in the presence of PC or PC/PA liposomes. Bars indicate mean ± SEM from n=3-4 independent experiments.* p<0.0001; Student’s t-test. **D.** Representative Western blots showing α-syn-140 and α-syn-112 binding to liposomes containing varying amounts of PA. Red box indicates liposome-containing fractions. α-Syn112 exhibited enhanced binding to PC/PA liposomes. **E.** Quantification of lipid bound protein. Data points indicate mean ± SEM from n=3-4 experiments. Asterisks indicate statistical significance using ANOVA (Sidak’s post hoc). * p<0.001. Data were best fit by a nonlinear dose-response curve (R^2^_(Syn140)_=0.96; R^2^_(Syn112)_=0.84).

We next investigated whether α-syn-112 also bound better to other anionic phospholipids, including PS and several of the phosphoinositides: phosphatidylinositol (PI), phosphoinositol 3-phosphate [PI(3)P], phosphoinositol 4-phosphate [PI(4)P], phosphoinositol 4,5-bisphosphate [PI(4,5)P_2_], and phosphoinositol 3,4,5-triphosphate [PI(3,4,5)P_3_]. PI(4)P and PI(4,5)P_2_ are of particular interest to this study because of their known roles in regulating synaptic vesicle exocytosis and endocytosis (Wenk et al., 2001; Di Paolo and De Camilli, 2006; Saheki and De Camilli, 2012). For these experiments, we used 5% of the anionic phospholipids because this was the sub-saturating condition with the greatest differential binding in the PS concentration series (Fig. 2). For all lipids tested, PS, PI, PI(3)P, PI(4)P, PI(4,5)P_2_, and PI(3,4,5)P_3_ alike, α-syn-112 bound significantly better than α-syn-140 (Fig. 3A). Quantification of band intensities from 3-6 replicates revealed that in most cases α-syn-112 binding to the anionic phospholipids was on average at least 2-fold greater than α-syn-140 (Fig. 3B) [(*PS*= α-syn-140: 34.42 ± 10.07, n=5; α-syn-112: 71.50 ± 2.76, n=6; p=0.0038; Student’s t-test) (*PI*= α-syn-140: 24.90 ± 10.73, n=3; α-syn-112: 71.78 ± 5.91, n=3; p=0.0187; Student’s t-test) (*PI(3)P*= α-syn-140: 22.83 ± 9.1, n=3; α-syn-112: 66.86 ± 3.78, n=4; p=0.0014; Student’s t-test) (*PI(4)P*= α-syn-140: 20.91 ± 6.05, n=3; α-syn-112: 57.07 ± 8.83, n=4; p=0.0264; Student’s t-test) *(PI(4,5)P_2_=* α-syn-140: 43.68 ± 3.00, n=4; α-syn-112: 67.50 ± 2.48, n=4; p=0.0009; Student’s t-test) (*PI(3,4,5)P_3_*= α-syn-140: 35.02 ± 1.457, n=3; α-syn-112: 87.84 ± 6.33, n=3; p=0.0012; Student’s t-test)]. Taken together, these data reveal that α-syn-112 binds significantly better than α-syn-140 to all negatively-charged anionic phospholipids tested.

**Figure 3.**
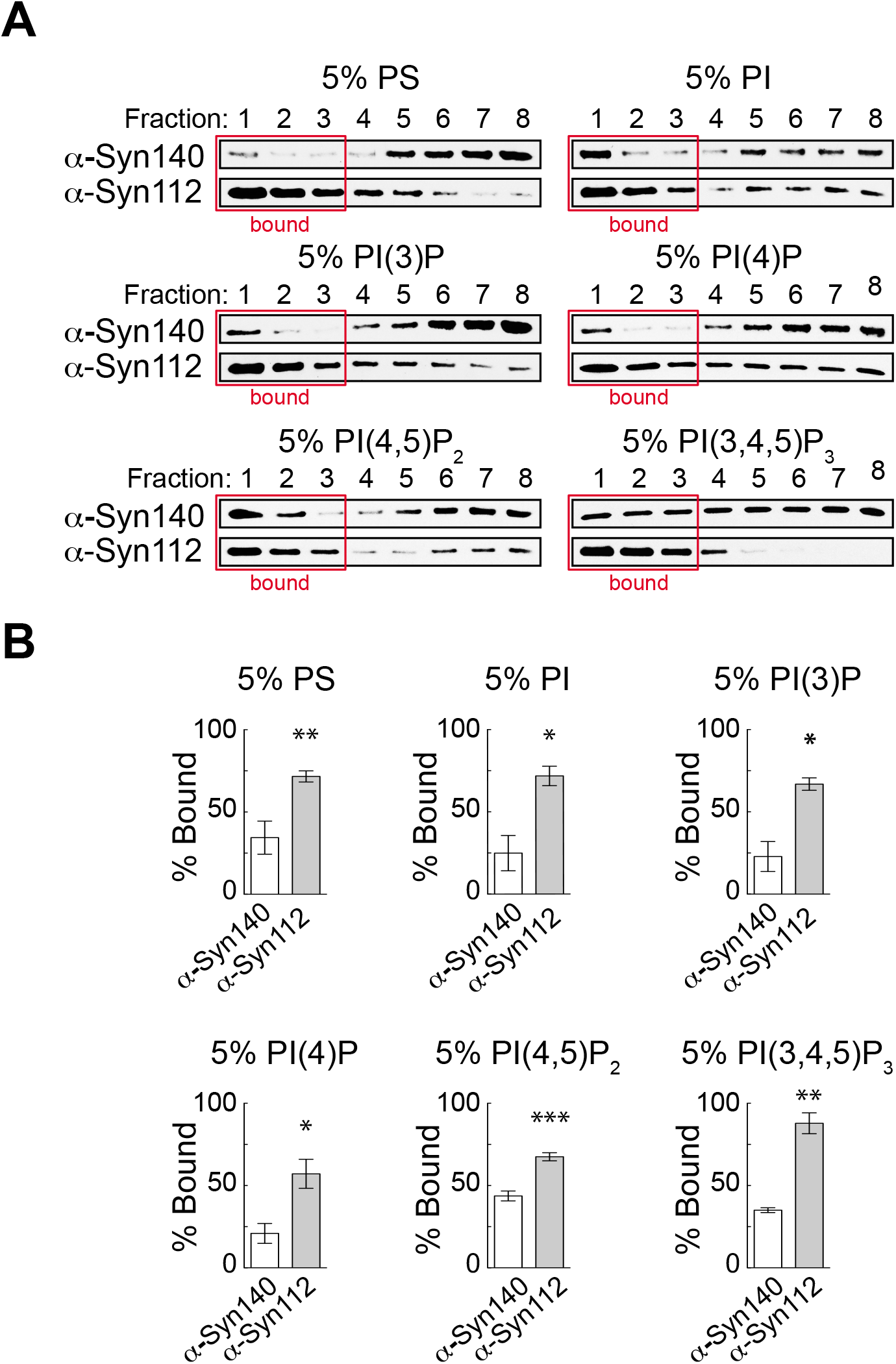
α -Syn-112 exhibits enhanced binding to anionic phospholipids. **A.** Representative Western blots showing α-syn-140 and α-syn-112 binding to liposomes containing 5% PS, PI, PI(3)P, PI(4)P, PI(4,5)P_2_ and PI(3,4,5)P_3_. Red box indicates liposome-containing fractions. α-Syn-112 bound with increased efficacy to all liposomes containing anionic phospholipids. **B.** Quantification of lipid bound protein show enhanced binding to all anionic phospholipids tested. Data points indicate mean ± SEM from n=3-6 experiments. Asterisks indicate statistical significance by Student’s t-test. * p<0.05, ** p<0.01, *** p<0.001.

### Enhanced binding of α-syn-112 to liposomes generated from total brain lipids

We next wanted to evaluate binding of α-syn-140 and α-syn-112 to a complex mixture of lipids that is more physiologically relevant. We therefore tested binding to liposomes made from purified brain total lipid extract (BTLE). HPLC analysis indicates that a substantial percentage of the BTLE lipids comprise PC, PA, PS, and PI, lipids we have already tested individually, as well ~59% unknown lipids (Fig. 4A). After ultracentrifugation in the Accudenz gradient, >80% of BTLE liposomes floated to the top of the column (Fig. 4B). As was observed with the individual anionic lipids, α-syn-112 exhibited a >2-fold greater binding to BTLE liposomes compared to α-syn-140 (Fig. 4C-D) (α-syn-140: 36.46 ± 11.56, n=3; α-syn-112: 87.21 ± 4.096, n=6; p=0.0144; Student’s t-test). Thus, α-syn-112 still binds significantly better than α-syn-140 when presented with a complex mixture of brain-derived liposomes.

**Figure 4.**
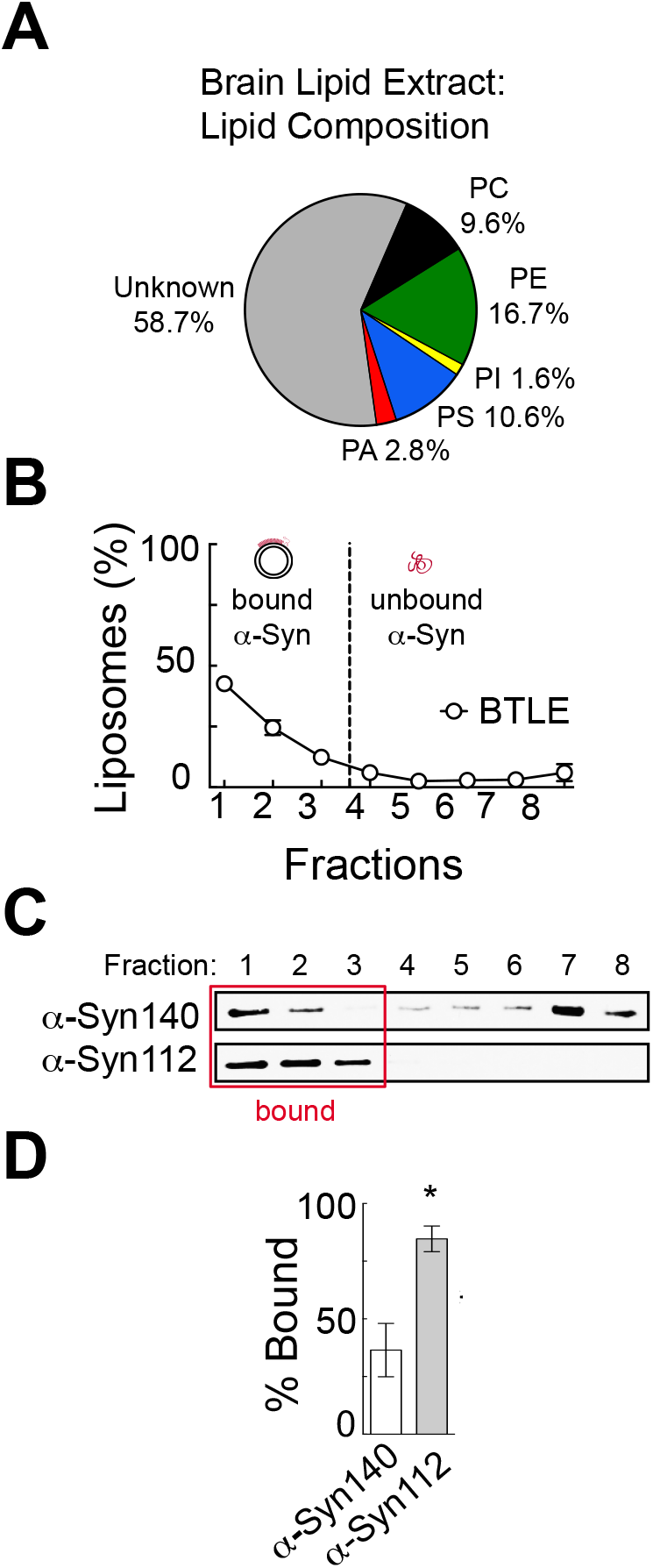
α -Syn-112 exhibits enhanced binding to total brain lipids. **A.** Pie chart showing the percentages of known phospholipids in brain total lipid extracts (BTLE) from porcine brain, which were used to make liposomes. **B.** Distribution of BTLE liposomes after ultracentrifugation. **C.** Representative Western blots showing α-syn-140 and α-syn-112 binding to liposomes. Red box indicates liposome-containing fractions. **D.** Quantification of lipid bound protein indicates that α-syn-112 has enhanced binding to BTLE liposomes. Bars indicate mean ± SEM from n=3 experiments. Asterisk indicates p=0.0144 by Student’s t-test.

### α-Syn-112 exhibits enhanced oligomerization on synaptic membranes

We next examined how α-syn-140 and α-syn-112 interact with physiological synaptic membranes using an established membrane recruitment assay (Shetty et al., 2013). First, synaptosome membranes were isolated from mouse brain and stripped of all associated proteins. Endogenous α-syn-140 was not detected on the stripped membranes, while transmembrane proteins such as N-cadherin were retained (Fig. 5A). The stripped membranes were then incubated with cytosolic proteins alone or supplemented with 2 μM recombinant human α-syn-140 or α-syn-112. Under these conditions, we observed robust recruitment of both isoforms to synaptic membranes, but with different patterns (Fig. 5B). α-Syn-140 was recruited predominantly in the monomeric form (76.6% ± 1.7, n=4), and its oligomerization into dimers and trimers comprised a small fraction (dimer: 22.6% ± 1.8; trimer: 0.79% ± 1.8; n=4). In contrast, after membrane recruitment, α-syn-112 was mostly dimeric and trimeric under the same conditions (monomer: 34.6 ± 5.8%; dimer: 56.4% ± 4.2; trimer: 9.0% ± 1.9; n=4), indicating that α-syn-112 is prone to oligomerization on synaptic membranes (Fig. 5B-C). This oligomerization was triggered by the interaction with synaptic membranes since both α-syn-140 and α-syn-112 were >90% monomeric in the starting material (Fig. 5B-C, left lanes) [(α-syn-140 - monomer: 92.9% ± 2.3; dimer: 5.7% ± 2.3; trimer: 1.3% ± 0.2; n=9) (α-syn-112 - monomer: 91.7% 5.8; dimer: 6.8% ± 2.9; trimer: 1.3% ± 0.3; n=9). Further demonstrating the enhanced oligomerization of α-syn-112, the oligomer/monomer ratio for α-syn-112 was 7-fold greater than that of α-syn-140 (Fig. 5D) (α-syn-140= 0.31 ± 0.03; α-syn-112= 2.16 ± 0.53, n=4; p=0.0067, Student’s t-test). When the membrane recruitment assay was repeated using varying concentrations of α-synuclein, α-syn-140 bound synaptic membranes primarily in the monomeric form with only ~30% oligomerization observed at the highest concentrations (Fig. 5E). In contrast, at all concentrations tested, where detectable, α-syn-112 bound to synaptic membranes as monomers, dimers, and trimers, with the proportion of oligomeric species exceeding that of monomeric α-syn-112 (Fig. 5F). Thus, α-syn-112 exhibits greater oligomerization on purified synaptic membranes.

**Figure 5.**
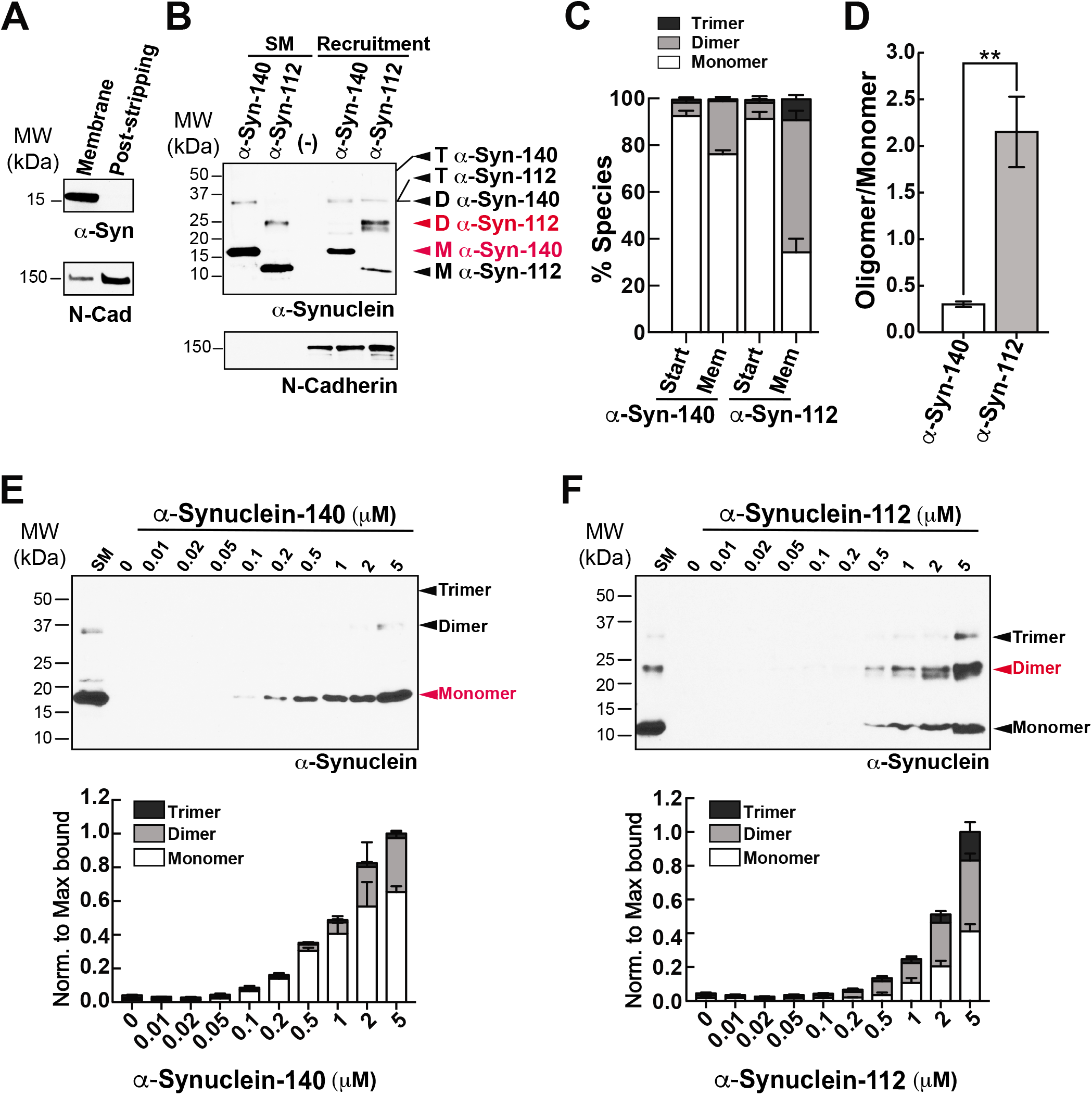
Enhanced oligomerization of α -syn-112 on synaptic membranes. **A.** Purified synaptic membranes from mouse brain are efficiently stripped of membrane associated proteins such as endogenous α-synuclein, while transmembrane proteins like N-cadherin remain. **B-C.** Recruitment of recombinant α-syn-140 and α-syn-112 to stripped synaptic membranes in the presence of brain cytosol. Compared to α-syn-140, which was recruited to synaptic membranes predominantly as a 15 kDa monomer (M), α-syn-112 bound predominantly as a 25 kD dimer (D) or trimer (T). SM = starting material. (-) indicates the cytosol only control. **D.** α-Syn-112 shows a 7-fold increase in the oligomer/monomer ratio compared to α-syn-140. Bars indicate mean ± SEM from n=3-4 experiments. Asterisks indicate statistical significance using Student’s t-test (** p<0.01). **E-F.** Concentration curve for α-syn-140 and α-syn-112 binding to synaptic membranes. α-Syn-112 oligomerized more at all concentrations tested. Red indicates the most abundant molecular species.

### Excess α-syn-112 impairs synaptic vesicle recycling, producing effects consistent with enhanced dimerization

We previously reported that acute introduction of α-syn-140 severely impaired synaptic vesicle endocytosis at lamprey synapses (Busch et al., 2014; Medeiros et al., 2017; Banks et al., 2020), a finding that was corroborated at mammalian synapses (Xu et al., 2016; Eguchi et al., 2017). In contrast, α-synuclein mutants with reduced membrane capacity, such as the point mutant A30P, produce little to no deficits in synaptic vesicle trafficking (Nemani et al., 2010; Busch et al., 2014), suggesting that lipid binding capacity is a strong predictor of the severity of synaptic defects. We therefore hypothesized that α-syn-112 would similarly induce synaptic vesicle recycling defects that were as or more robust than those reported for α-syn-140. To test this, giant RS axons were microinjected with recombinant human α-syn-112 (120-200 μM pipet concentration), as was done for α-syn-140 in our prior studies (Busch et al., 2014; Medeiros et al., 2017; Banks et al., 2020), thus directly delivering the protein to the presynapses (Walsh et al., 2018). Upon axonal injection, the protein is diluted 10-20x for a final concentration of 7-16 μM, which is 2-3 times the estimated endogenous levels of α-synuclein at mammalian synapses (Westphal and Chandra, 2013) and consistent with overexpression levels observed in PD (Singleton et al., 2003). After injection, axons were stimulated (20 Hz, 5 min), fixed and processed for standard transmission electron microscopy, as previously described (Busch et al., 2014; Medeiros et al., 2017; Walsh et al., 2018). Images of control synapses were obtained from the α-synuclein-injected axons but at distances beyond where the protein diffused (based on a co-injected fluorescent dye), thus providing an internal control for each experiment.

Giant RS synapses are *en passant* glutamatergic synapses that reside along the perimeter of the giant RS axons (Wickelgren et al., 1985; Brodin and Shupliakov, 2006). Stimulated control synapses exhibit a large and localized synaptic vesicle cluster, shallow plasma membrane evaginations, few clathrin-coated pits (CCPs) and clathrin-coated vesicles (CCVs), and only a few cisternae, which are defined as large vesicular structures with a diameter >100 μm (Fig. 6A). While we do not yet know the precise identities of cisternae, their morphologies are consistent with bulk and/or recycling endosomes (Morgan et al., 2013; Chanaday et al., 2019). By comparison, synapses treated with recombinant human α-syn-112 exhibited a drastic change in morphology, indicated by a loss of the synaptic vesicle cluster, large extended plasma membrane evaginations, and accumulation of cisternae and clathrin-coated pits and vesicles (Fig. 6B). Three-dimensional reconstructions show clearly the morphological alterations caused by α*-*syn-112, especially the loss of vesicles (blue) and buildup of plasma membrane (green) and cisternae (magenta) (Fig. 6C-D). In addition, there were obvious changes in the number of CCPs and CCVs. Whereas stimulated control synapses have only a few CCPs, those treated with α-syn-112 have more CCPs and CCVs, suggesting deficits in vesicle fission and clathrin uncoating (Fig. 6E-G). A quantitative analysis revealed that introduction of excess α-syn-112 significantly reduced the average number of synaptic vesicles per synapse (per section) by almost 70% (Fig. 6H) (Control: 133.2 ± 12.94 SVs, n=25 synapses, 2 axons/animals; α-syn-112: 42.58 ± 4.175 SVs, n=24 synapses, 2 axons/animals; p<0.0001; Student’s t-test). The remaining synaptic vesicles had larger diameters (Control: 50.6 ± 0.5 nm, n=200 SVs, 5 synapses; α-syn-112: 53.0 ± 0.9 nm, n=200 SVs, 5 synapses; p<0.0001; Student’s t-test). The loss of synaptic vesicles was compensated by a significant increase in the size of the plasma membrane evaginations, indicating a defect in synaptic vesicle endocytosis (Fig. 6I) (Control: 2.417 ± 0.1898, n=25; α-syn-112: 3.951 ± 0.1947, n=24; p<0.0001; Student’s t-test). Consistent with endocytic defects, the number and size of cisternae were also increased (Fig. 6J-K) [(*# Cisternae:* Control: 2.440 ± 0.3655, n=25; α-syn-112: 8.167 ± 1.324, n=24; p=0.0001; Student’s t-test) *(Cisternae Size:* Control: 0.3830 ± 0.01850, n=68; α-syn-112: 0.4908 ± 0.02044, n=196; p=0.0033; Student’s T-test)]. The total number of combined CCP/Vs also increased more than 2-fold (Fig. 6L) (Control: 1.680 ± 1.249, n=25; α-syn-112: 3.417 ± 2.376, n=24; p=0.0023; Student’s t-test). Analysis of the progressive stages of clathrin-mediated endocytosis revealed that α-syn-112 induced more constricted CCPs (stage 3) and free CCVs (stage 4), indicating that both fission and clathrin uncoating were impaired (Fig. 6M) *(Stage 1* Control: 0.1200 ± 0.3317 CCPs/section/synapse (n=25), α-Syn-112: 0.08333 ± 0.2823 CCPs/section/synapse (n=24); *Stage 2* Control: 0.2800 ± 0.4583 CCPs (n=25), α-Syn-112: 0.1250 ± 0.3378 CCPs (n=24); *Stage 3* Control: 0.9600 ± 0.9345 CCPs (n=25), α-Syn-112: 1.667 ± 1.435 CCPs (n=24); p<0.0001; ANOVA Sidak’s Post Hoc; *Stage 4* Control: 0.3200 ± 0.6904 CCVs (n=25), α-Syn-112: 1.542 ± 1.474 CCVs (n=24); p<0.0001; ANOVA Sidak’s Post Hoc). A total membrane analysis indicates that synaptic vesicle membrane was redistributed to plasma membrane, cisternae, and CCP/Vs in α-syn-112 treated synapses (Fig. 6N). These EM data indicate that like α-syn-140, α-syn-112 robustly impairs synaptic vesicle recycling consistent with effects on clathrin-mediated endocytosis and possibly bulk endocytosis (Busch et al., 2014; Medeiros et al., 2017; Medeiros et al., 2018; Banks et al., 2020).

**Figure 6.**
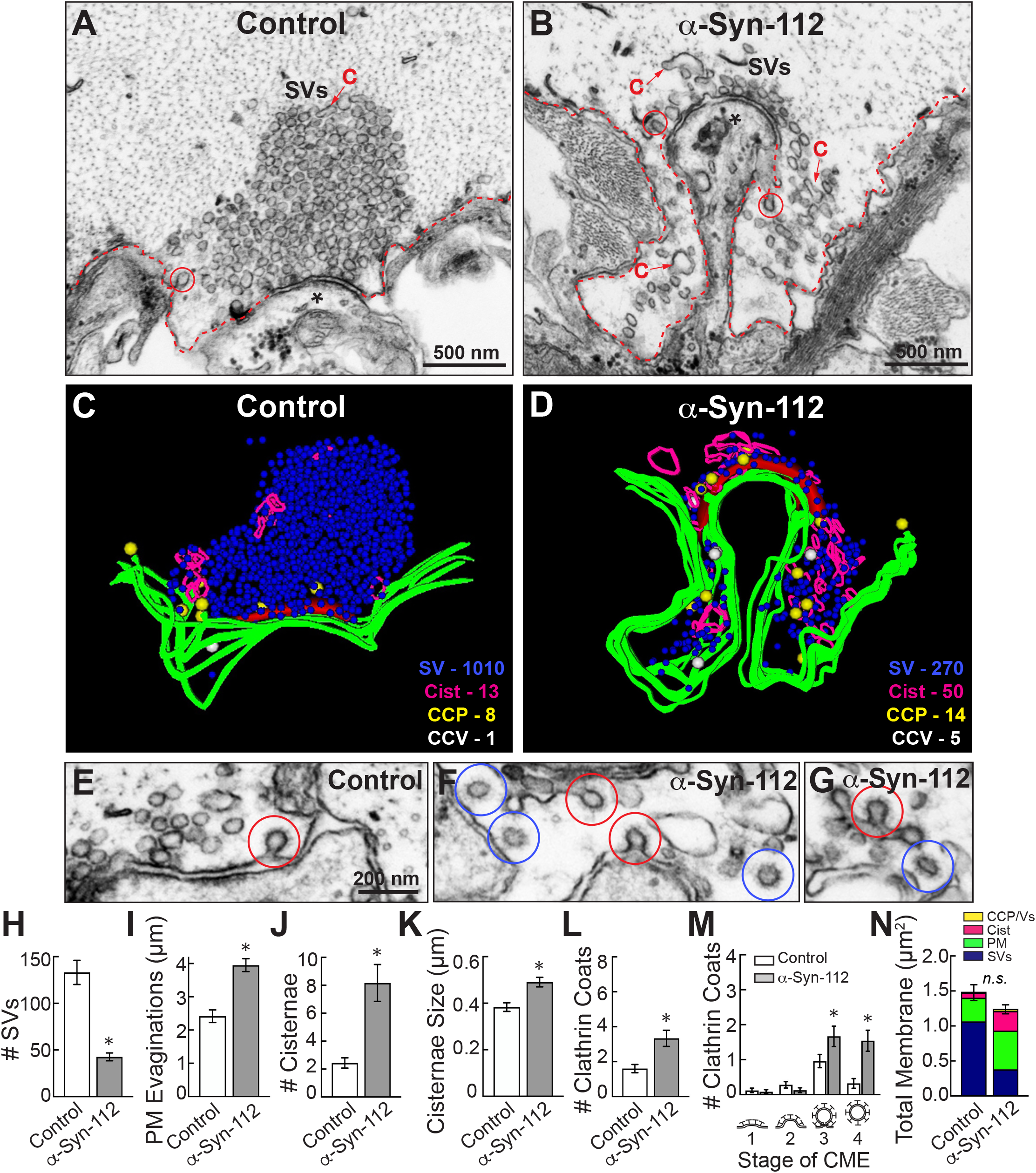
α -Syn-112 impairs synaptic vesicle recycling. **A-B.** Electron micrographs showing a stimulated (20 Hz, 5 min) control synapse and a synapse treated with recombinant human α-syn-112. While control synapses have a large pool of synaptic vesicles (SVs), moderate plasma membrane (PM) evaginations (red dotted line), and few cisternae (C) or clathrin coated structures (circles), those treated with α-syn-112 are dramatically altered with a notable loss of SVs and buildup of PM. Asterisk marks the postsynaptic dendrite. **C-D.** 3D reconstructions reveal that excess α-syn-112 induces a substantial loss of SVs, which was compensated by an extensive build up of plasma membrane (green), as well as increased numbers of cisternae (magenta) and clathrin-coated pits (CCPs; yellow) and clathrin-coated vesicles (CCVs; white), indicating impaired vesicle endocytosis. **E-G.** Micrographs showing effects of α-syn-112 on clathrin-mediated endocytosis. Control synapses have only few CCPs (red circles), while α-syn-112 treated synapses have many CCPs and CCVs (blue circles). Scale bar in E applies to F-G. **H-M.** Morphometric analyses showing differences between control and α-syn-112 treated synapses, which are consistent with defects in synaptic vesicle endocytosis. Increase in stage 3 CCPs and stage 4 CCVs indicates impaired vesicle fission and clathrin uncoating, respectively. Bars indicate mean ± SEM from n=25-26 synapses, 2 axons/animals. Asterisks indicate p<0.05 by Student’s t-test. **N.** Total membrane analysis shows redistribution of synaptic membranes by α-syn-112. n.s. = not significant.

While the synaptic phenotype produced by α-syn-112 overlaps with that previously reported for α-syn-140 (Busch et al., 2014; Medeiros et al., 2017; Banks et al., 2020), we also noted some distinct differences. Monomeric α-syn-140 induces CCV uncoating defects with no effect on earlier stages of CCP formation, while dimeric α-syn-140 primarily impairs CCP fission (Medeiros et al., 2017; Medeiros et al., 2018). In comparison, α-syn-112, which was initially injected in the monomeric form (see Fig. 1C), induced both CCP fission and CCV uncoating defects, demonstrated by the increase in stage 3 and 4 clathrin coats, suggesting some dimerization *in vivo* (Fig. 6M). In addition, α-syn-112 induced atypically deep plasma membrane evaginations around the active zone (Fig. 7A-D). These plasma membrane evaginations appeared deeper than those at α-syn-140 treated synapses (Fig. 7E-F) and more similar to those produced by dimeric α-syn-140 (Fig. 7G-H) (Medeiros et al., 2017). To quantify this effect, we measured the depth of the plasma membrane evaginations from the axolemmal surface to the deepest point within each evagination. Compared to control synapses, the plasma membrane evaginations produced by α-syn-112 were ~2-fold deeper (Fig. 7I), and they were quantitatively intermediate to those induced by monomeric and dimeric α-syn-140 (Fig. 7I) (Control: 1.00 ± 0.07, n=79 synapses, 2 axons/animals; α-Syn-112: 1.97 ± 0.12, n=24 synapses, 2 axons/animals; α-Syn-140: 1.49 ± 0.16, n=26 synapses, 2 axons/animals; α-Syn-140 Dimer: 2.40 ± 0.21, n=21,2 axons/animals; ANOVA, p<0.0001). Thus, consistent with its *in vitro* biochemical properties, α-syn-112 also produced striking phenotypes on synaptic membranes *in vivo* that are consistent with enhanced membrane binding and oligomerization.

**Figure 7.**
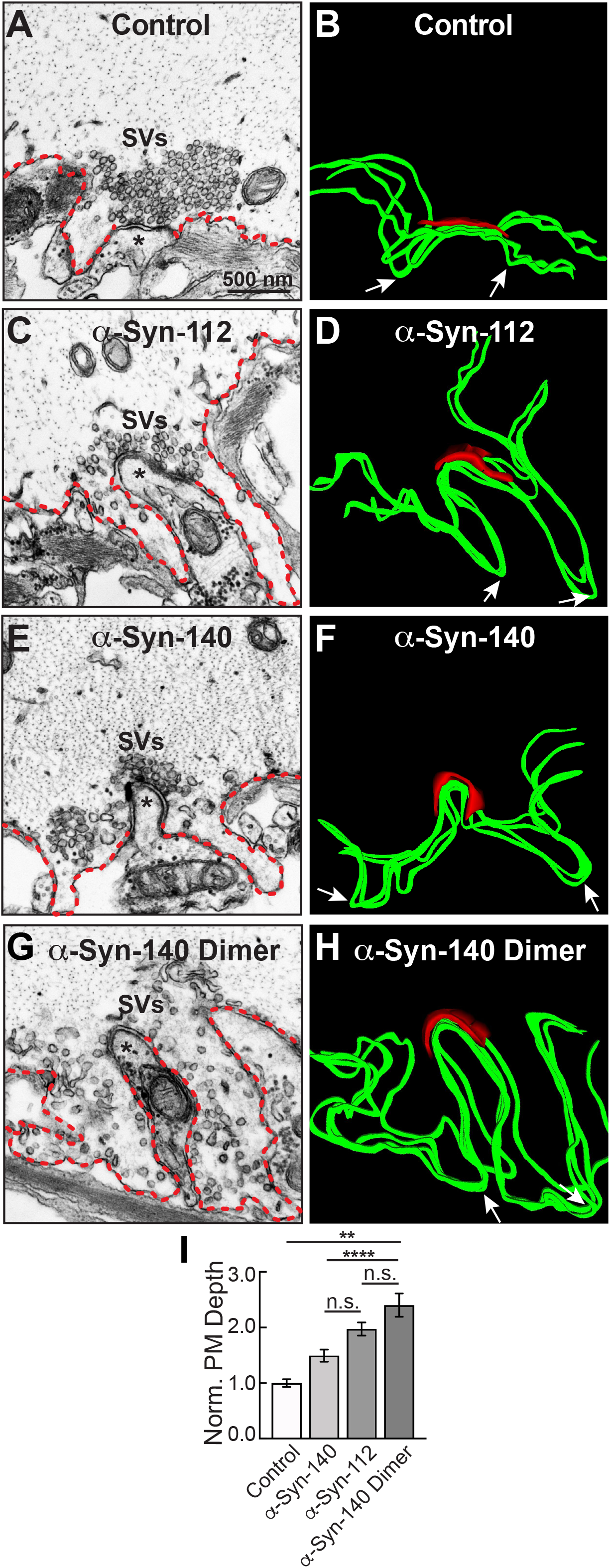
α -Syn-112 induces large plasma membrane evaginations intermediate to monomeric and dimeric α -syn-140. **A-B.** Electron micrograph (left) and 3D reconstruction of plasma membrane (right) at a control synapse. PM evaginations are small (red dotted line in micrograph; green ribbon in 3D reconstruction). Arrows in B, D, F and H indicate the deepest point in each evagination. Asterisks in A, C, E and G indicate the postsynaptic dendrite. **C-D.** In comparison to controls, acute introduction of α-syn-112 induced PM evaginations that were much larger and deeper. **E-H.** Plasma membrane evaginations produced by monomeric α-syn-140 (E-F) and dimeric α-syn-140 (G-H) were also enlarged compared to controls. **I.** Quantification of the depth of PM evaginations reveals that α-syn-112 has an intermediary phenotype to monomeric and dimeric α-syn-140, consistent with its ability to dimerize on synaptic membranes. Bars indicate mean ± SEM from n=21-26 synapses, 2 axons/animals. Asterisks indicate statistical significance by one-way ANOVA ** (p<0.01); **** (p<0.0001); n.s.=not significant.

## DISCUSSION

This is the first study to investigate the lipid binding properties and synaptic effects of α-syn-112, which is both naturally occurring and overexpressed in multiple neurodegenerative diseases. We show here that α-syn-112 exhibits enhanced membrane binding *in vitro* compared to wild type α-syn-140 (Fig. 2–4), including to synaptically relevant phosphoinositides such as PI(4)P and PI(4,5)P_2_ (Fig. 3). α-Syn-112 also exhibits enhanced oligomerization (dimerization and trimerization) on synaptic membranes (Fig. 5) and impairs synaptic vesicle recycling when acutely introduced in excess (Fig. 6–7). In our previous studies, we showed that excess monomeric α-syn-140 impaired CCV uncoating at lamprey synapses (Medeiros et al., 2017; Banks et al., 2020), while dimeric α-syn-140 impaired an earlier stage of CCP fission (Medeiros et al., 2017; Medeiros et al., 2018). Interestingly, although we injected recombinant α-syn-112 in the monomeric form (Fig. 1C), the resulting synaptic phenotype was indicative of deficits in both CCP fission and CCV uncoating (Fig. 6), which is consistent with its enhanced ability to dimerize on synaptic membranes (Fig. 5). Further underscoring this result is that the depth of plasma membrane evaginations produced by α-syn-112 was also intermediate between monomeric and dimeric α-syn-140. We do not yet fully understand the oligomerization status of α-syn-112 once it enters the synaptic environment. However, this study nonetheless further emphasizes that different α-synuclein species can product distinct effects at synapses (Medeiros et al, 2018), which may compound the cellular deficits if expressed combinatorially.

A key biochemical feature of α-syn-112 is its ability to bind phospholipid membranes with increased efficacy, as compared to wild type α-syn-140. In every example tested, α-syn-112 exhibited enhanced binding *in vitro* to anionic phospholipids, including many of the phosphoinositides and total brain lipids (Fig. 2–4). The predicted structure for α-syn-112 involves a deletion of 28 amino acids (a.a. 103-130) in the C-terminal domain, which may result in an extended alpha helical region (Fig. 1B). Given that the membrane binding capacity of α-syn-140 is fairly evenly distributed throughout the alpha helical N-terminal domain (Davidson et al., 1998; Chandra et al., 2003; Burre et al., 2012), extending the alpha helix could result in the enhanced lipid binding that was observed. Additionally, we show that α-syn-112 also binds more strongly to a number of phosphoinositides, including PI, PI(3)P, PI(4)P, PI(4,5)P_2_, and PI(3,4,5)P_3_, though we did not detect any preferential selectivity amongst them (Fig. 3). While interactions between α-syn-140 and PI(4,5)P_2_ have been reported using giant unilamellar vesicles (Narayanan et al., 2005; Stockl et al., 2008), to our knowledge this is the first study that provides a more comprehensive and comparative assessment of α-synuclein binding to phosphoinositides. It is notable that such strong binding was observed when the phosphoinositide concentrations were only 5% of the total lipid composition (Fig. 2), which is much less than the 30-50% anionic lipids normally used in these *in vitro* assays (Burre et al., 2012; Busch et al., 2014; Medeiros et al., 2017). Phosphoinositides are present in limiting amounts and tightly-controlled on cellular membranes (Di Paolo and De Camilli, 2006; Takamori et al., 2006; Balla, 2013; Schink et al., 2016), including on synaptic vesicles where they likely comprise <10% of the total phospholipids (Takamori et al., 2006). Thus our results may be more reflective of what happens intracellularly and suggest that α-syn-112 attaches to physiological synaptic membranes better than α-syn-140, which has implications for its potential toxicity.

PI(4,5)P_2_ is enriched on the plasma membrane and helps to recruit clathrin adaptor proteins to the membrane during initiation of clathrin-mediated synaptic vesicle endocytosis (Ford et al., 2001; Di Paolo and De Camilli, 2006; Saheki and De Camilli, 2012). Thus, the strong binding of α-syn-140 and α-syn-112 to PI(4,5)P_2_ may mask sites for clathrin coat initiation and inhibit early stages of vesicle endocytosis, which is consistent with the expanded plasma membrane evaginations observed after introducing either isoform to synapses (Fig. 6–7) (Busch and Morgan, 2012; Medeiros et al., 2017; Banks et al., 2020). Stronger binding to PI(4,5)P_2_ may also explain in part why α-syn-112 has greater effects than α-syn-140 on the depth of plasma membrane evaginations (Fig. 7). In addition, it is thought that PI(4,5)P_2_ remains on the endocytic vesicle throughout CCP and CCV formation until it is dephosphorylated to PI(4)P by the uncoating protein, synaptojanin (Cremona et al., 1999; Saheki and De Camilli, 2012). Thus, strong binding of α-syn-140 and α-syn-112 to PI(4,5)P_2_ and PI(4)P may also mask these lipids and alter the dynamics of the late stages of clathrin-mediated endocytosis and contribute to the fission and uncoating defects observed, along with mislocalization of the CCV uncoating protein (Hsc70), which we recently reported (Banks et al., 2020). Going forward, it will be important to advance our understanding of α-synuclein interactions with phosophoinositides, since misregulation of phosphoinositide levels and phosphoinositide-mediated membrane trafficking may contribute to neurodegenerative diseases (Fabelo et al., 2011; Nadiminti et al., 2018)

Another interesting finding is that α-syn112 has increased propensity for oligomerization on synaptic membranes (Fig. 5). Like monomeric α-syn-140, dimeric α-syn-140 undergoes alpha helical folding in the presence of SDS micelles, binds strongly to PA-containing liposomes, and exhibits time-dependent aggregation and fibrillation *in vitro* in biochemical assays (Pivato et al., 2012; Medeiros et al., 2017; Dong et al., 2018). α-Synuclein rapidly dimerizes and aggregates on membranes containing PS (Lv et al., 2019), which is one of the major anionic lipids comprising synaptic vesicles (Takamori et al., 2006). Under physiologic conditions, α-syn-140 multimers exist at synapses and participate in synaptic vesicle clustering, restricting vesicle motility during trafficking (Wang et al., 2014). When introduced in excess to synapses, dimeric α-syn-140 inhibited synaptic vesicle recycling and impaired CCP fission (Medeiros et al., 2017; Medeiros et al., 2018). Because excess α-syn-112 also interfered with CCP fission (Fig. 6), this suggests that the injected recombinant monomeric α-syn-112 protein dimerized upon interaction with synaptic membranes *in vivo,* consistent with its *in vitro* effects (Fig. 5). In future experiments, it will be interesting to determine the impacts of dimeric α-syn-112 on synaptic vesicle trafficking. Since oligomerization on membranes is associated with membrane penetration and toxicity (Tsigelny et al., 2012; Tsigelny et al., 2015), formation of α-syn-112 or α-syn-140 dimers may be an important rate-limiting step in the early pathogenesis of the synucleinopathies.

In summary, like α-syn-140, α-syn-112 avidly binds phospholipid membranes and, when in excess, impairs synaptic vesicle recycling producing distinct effects on clathrin-mediated endocytosis. Despite these similarities, α-syn-112’s enhanced membrane binding properties and propensity for oligomerization may underlie the greater effects on synaptic membranes. In addition to providing the first insight into the synaptic toxicity caused by α-syn-112, this study further emphasizes the need for investigating the impacts of different α-synuclein isoforms and conformations on neuronal function, since doing so may help us better understand the cellular pathways leading to neurodegeneration.

## ACKNOWLEDGMENTS

The authors would like to thank Louie Kerr from the Central Microscopy Facility at the Marine Biological Laboratory in Woods Hole, MA for technical support with electron microscopy. We also thank Dr. Luigi Bubacco (Univ. of Padova) for providing the recombinant α-synuclein dimer and Drs. Elizabeth Jonas and Timothy Eisen for critical feedback on the manuscript.

## AUTHOR CONTRIBUTIONS

Authors LS, JE, KV, AM and JM contributed to the conception and design of the study, as well as data acquisition, analysis and interpretation. KH contributed to data acquisition. All authors (LS, JE, KV, AM, KH, JM) were involved in drafting the manuscript, have provided final approval of this manuscript for submission, and agree to be accountable for all aspects of the work.

